# Batch correction methods used in single cell RNA-sequencing analyses are often poorly calibrated

**DOI:** 10.1101/2024.03.19.585562

**Authors:** Sindri Emmanúel Antonsson, Páll Melsted

**Affiliations:** Faculty of Industrial Engineering, Mechanical Engineering, and Computer Science, University of Iceland, Reykjavík, Iceland

## Abstract

As the number of experiments that employ single-cell RNA-sequencing (scRNA-seq) grows it opens up the possibility of combining results across experiments or processing cells from the same experiment assayed in separate sequencing runs. The gain in the number of cells that can be compared comes at the cost of batch effects that may be present. Several methods have been proposed to combat this for scRNA-seq datasets.

We compared seven widely used method used for batch correction of scRNA-seq datasets. We present a novel approach to measure the degree to which the methods alter the data in the process of batch correction, both at the fine scale comparing distances between cells as well as measuring effects observed across clusters of cells. We demonstrate that many of the published method are poorly calibrated in the sense that the process of correction creates measurable artifacts in the data.

In particular, MNN, SCVI and LIGER performed poorly in our tests, often altering the data considerably. Batch correction with Combat, BBKNN and Seurat introduced artifacts that could be detected in our setup. However, we found that Harmony was the only method that consistently performed well, in all the testing methodology we present. Due to these result Harmony is the only method we can safely recommend using when performing batch correction of scRNA-seq data.

## Introduction

With the increasing complexity and scale of scRNA-seq experiments, the need to integrate and combine differently collected data has also increased. Large scale datasets enable researchers to combine and compare different data modalities to gain increased insight and accuracy. When scRNA-seq data are collected at different times, with different protocols, technologies or sequencing platforms, the integeration becomes increasingly complex. All these factors can affect the expressions of genes in complex ways, some of these differences can be biological in origin, others from technical artifacts. We aggregate the varation due to technical artifacts under the umbrella term of batch effects.

There are some unique challenges in integrating batches of scRNA-seq data that are not present when working with bulk RNA-seq data. Cell type composition can differ between batches and within cell types there can be systematic differences in gene expression between batches (Luecken and Theis 2019). One of the first steps in processing scRNA-seq data is to cluster or identify cells by cell type, thus requiring batch correction methods tailored for scRNA-seq datasets in order to ensure that cells of the same type are grouped together across batches.

Other studies have evaluated the performance of batch correction methods, focusing more on measuring how well batch effects are removed rather than how well the variation of interest is retained or whether the methods are well calibrated. Early work (Tran et al. 2020) compared 14 different batch correction methods based on running time, and classical metrics from cluster comparison. A recent preprint (Tyler et al. 2021) approached the problem of batch correction from both viewpoints, evaluating how well batch effects are corrected and the extent to which real biological effects are preserved after batch correction.

Any systematic effects on gene expression, affecting a large number of genes, will affect each point of the computational pipeline that starts with the raw sequencing data or count matrix and ends with a statistical test that is computed in order to demonstrate a biological difference. This systematic bias must be addressed, regardless of whether the effect is biological in origin, e.g. cells from different environments or individuals, or technical, i.e. the result of batch effects.

While most of the proposed methods have demonstrated, in their respective manuscripts, that they are capable of removing batch effects, they do not show that in the absence of batch effects the batch “corrections” do not alter the underlying truth.

We propose a methodology to assess the quality of these methods and to measure the extent to which these methods alter the data in the process of batch correction, in particular the case when there is little or no batch effect. We perform the batch correction and examine the change, if any, the process of each batch correction method has on the data and all downstream results.

Ideally, the application of batch effect correction should not correct the data at all as measured by a statistical test, i.e. the methods should be well calibrated. Under this null hypothesis, any significant change can be classified as an artifact of batch correction.

We present different datasets and use them to establish a ground truth: the distances between cells and the KNN structure etc. We then split them into two equal parts to create pseudo batches.

We examine the effects of the batch correction on three representative stages of most common computational pipelines in scRNA-seq analysis: a. the k-nearest neighbor(KNN) graph computed from the count matrix, b. clustering and cell type identification, c. differential expression analysis between clusters. We evaluated seven computational methods for batch correction that are commonly used in scRNA-seq pipelines.

## Results

### Batch correction methods

We selected seven different batch correction methods for evaluation, based on how commonly they are used and the difference in their respective methodologies. All of the methods are available as stand alone libraries or part of a larger package in Python or R.

Each method approaches the problem of batch correction in a different way. While the algorithms and methods in the original publications follow various different workflows, we have identified common themes and shared strategies, between the methods, that are highlighted in Table 1. The methods differ, not only in how the correction is performed, but which data object is being corrected. Namely, some methods correct the count matrix directly, whereas others leave the original matrix intact and correct either the k-NN graph or an embedding used to compute the k-NN graph.

**Table 1:**
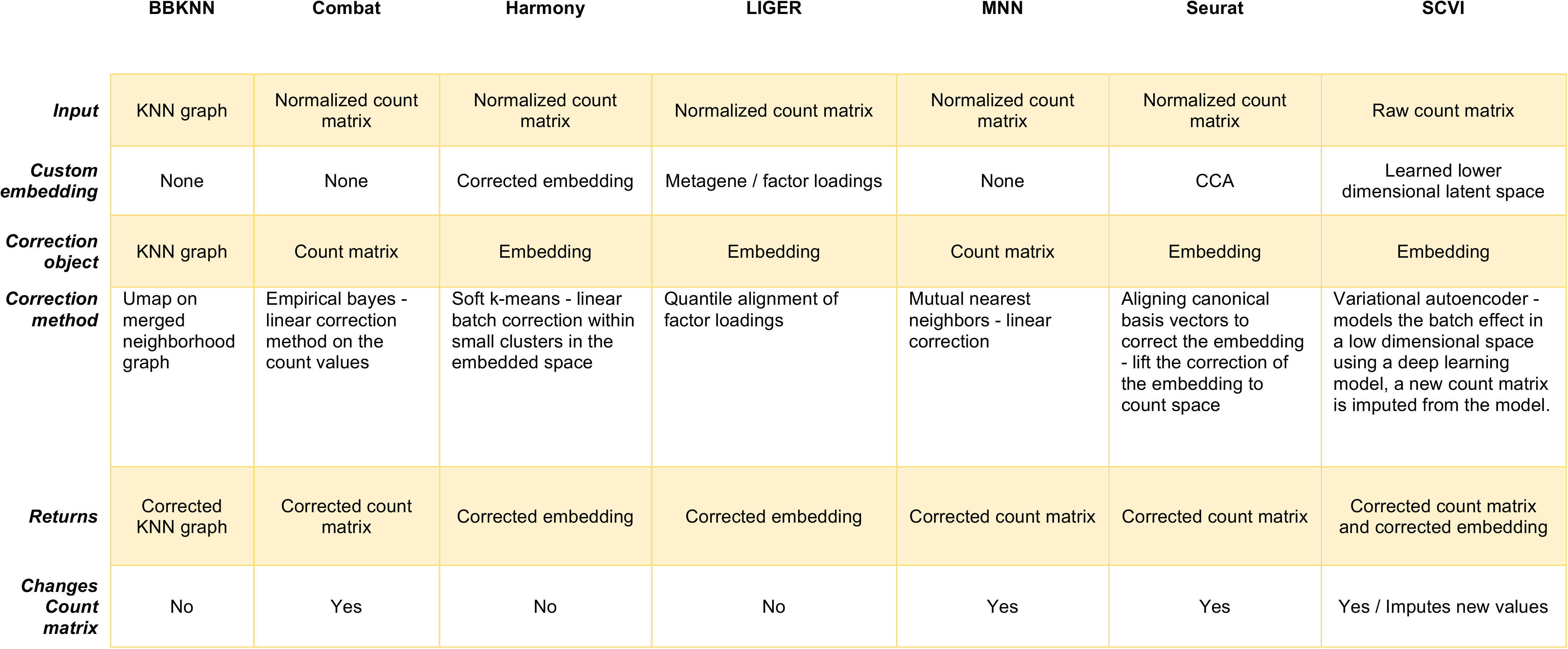
*Input* is the type of data that that particular method uses as input. The method may perform additional preprocessing steps on the input object before any calculations are performed. *Custom embedding* is the particular lower level embedding, if any, which the data is projected onto. The *correction object* is the actual data object that the method uses to make corrections. The *correction method* is a informal description of the particular method used for batch correction. *Returns* is the type of object the method returns. *Changes count matrix* is whether the method edits or returns a new count matrix to be used insted of the uncorrected count matrix in any subsequent steps in the workflow.

|  | BBKNN | Combat | Harmony | LIGER | MNN | Seurat | SCVI |
| --- | --- | --- | --- | --- | --- | --- | --- |
| **Input** | KNN graph | Normalized count matrix | Normalized count matrix | Normalized count matrix | Normalized count matrix | Normalized count matrix | Raw count matrix |
| **Custom embedding** | None | None | Corrected embedding | Metagene / factor loadings | None | CCA | Learned lower dimensional latent space |
| **Correction object** | KNN graph | Count matrix | Embedding | Embedding | Count matrix | Embedding | Embedding |
| **Correction method** | Umap on merged neighborhood graph | Empirical bayes - linear correction method on the count values | Soft k-means - linear batch correction within small clusters in the embedded space | Quantile alignment of factor loadings | Mutual nearest neighbors - linear correction | Aligning canonical basis vectors to correct the embedding - lift the correction of the embedding to count space | Variational autoencoder - models the batch effect in a low dimensional space using a deep learning model, a new count matrix is imputed from the model. |
| **Returns** | Corrected KNN graph | Corrected count matrix | Corrected embedding | Corrected embedding | Corrected count matrix | Corrected count matrix | Corrected count matrix and corrected embedding |
| **Changes Count matrix** | No | Yes | No | No | Yes | Yes | Yes / Imputes new values |

We measure the change caused by batch correction in: the count matrix, the low dimensional embedding of the count matrix, clustering and differential expression. Not all the methods alter the data at the same stage. Combat (Johnson et al. 2007), MNN (Haghverdi et al. 2018) and Seurat (Stuart et al. 2019) alter the count matrix, thus we can compare the altered count matrix to the ground truth and on any downstream results. Harmony(Korsunsky et al. 2019) corrects for batch effects by computing a low dimensional PCA(Principal component analysis) embedding and altering the embedding of each cell with respect to batches. BBKNN (Polański et al. 2020) changes only the k-NN graph based batch information. SCVI (Lopez et al. 2018) learns a low dimensional embedding of the data using a Variational Autoencoder in a Deep Learning framework, from which the corrected count matrices can be imputed. LIGER (Welch et al. 2019) creates a lower dimensional embedding and a combined KNN graph, which is then used to modify the lower dimensional embedding to correct for batch effects.

As Harmony, BBKNN, and LIGER do not modify the count matrix, the batch correction affects mainly the k-NN graph and all downstream results that rely on the k-NN graph, e.g. clustering. The expression estimates, in the count matrix, must then be corrected for batch effects using classical methods such as covariates in a linear model. In our results we have excluded these methods from comparisons where the explicit differences in the count matrix are computed.

### Simulation Strategy

The simplest simulation we evaluate is to generate a batch label where there is none. Namely, we take public scRNA-seq dataset and randomly label each cell as coming from batch A or batch B. This information is then supplied to each method which in turn corrects the dataset. If the batch correction methods are well calibrated we expect them to only make minor corrections that will not influence downstream analysis to a statistically significant degree.

Rather than simulating both count matrices and the random split, we use publicly available datasets on PBMCs from Humans and Neural cells from Mouse (Methods). The dataset were processed using standard preprocessing workflows (Methods) to obtain the cell by gene count matrix.

To do this we created random pseudo batches for each dataset, repeated 25 times (Figure 1A). For each iteration we perform the standard preprocessing (Methods) for the unaltered data, and randomly split the cells into 2 batches of equal size. We then apply the batch correction methods to the uncorrected count matrices with the pseudo batches as input. We then proceed with with the workflow, computing principle components, the k-NN graph, Leiden clustering and differential expression (Figure 1B, Methods). For each step we compare the computed artifact to their companion in the uncorrected data and report median values across the 25 iterations.

**Figure 1.**
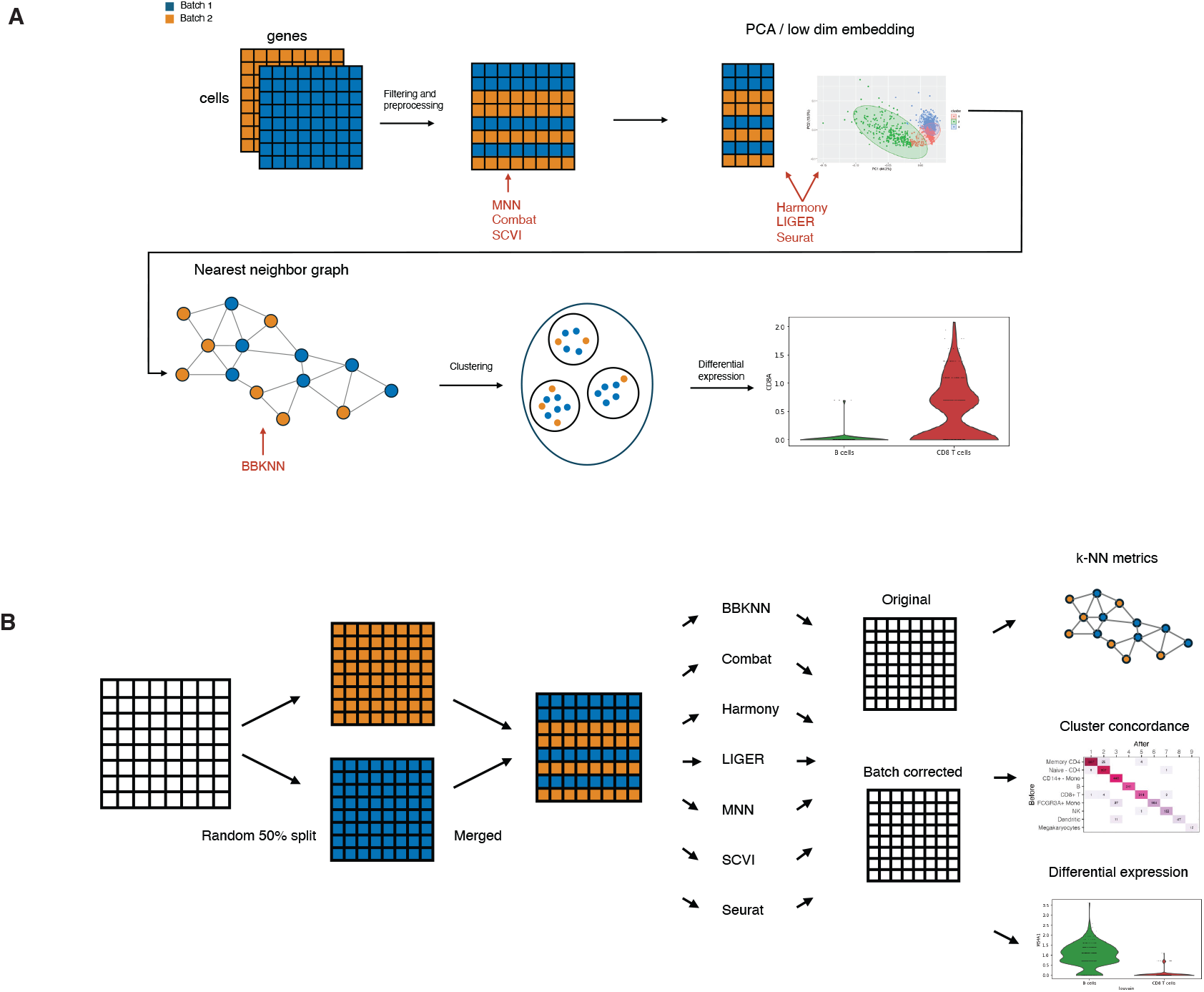
**A**: The fundamental steps of the batch correction workflow. Data is ingested in samples, combined and preprocessed in various ways, depending on the method. The data is then commonly projected onto a custom embedded space, possibly with lower dimension than the original data, where some correction takes place, correction can also be performed on the original count data or k-NN graph. The corrected object is then returned, depending on the method, a count matrix, lower dimensional embedding, NN graph etc. Downstream methods then use this corrected object. **B**: The general workflow of the evaluation conducted in this paper. SC rna-seq is split randomly into 2 batches. Batch correction methods are applied to these 2 pseudo batches. The change in the data, at different stages of the scRNA-seq workflow pipeline, is then measured to assess the change the act of batch correction has had on the data.

### Changes in cell type clusters

The batch corrections performed can influence not only gene expression but the cluster assignment of each cell. To investigate the effects we annotated the clusters of the uncorrected dataset (Methods, Supplement) and then matched the clusters obtained after batch correction. Figure 2 shows the cluster confusion matrix and the imbalance of batches within clusters for each correction method. In many cases the corrected data was split into more clusters, in those cases we manually ordered the clusters to make the display as diagonal as possible.

**Figure 2:**
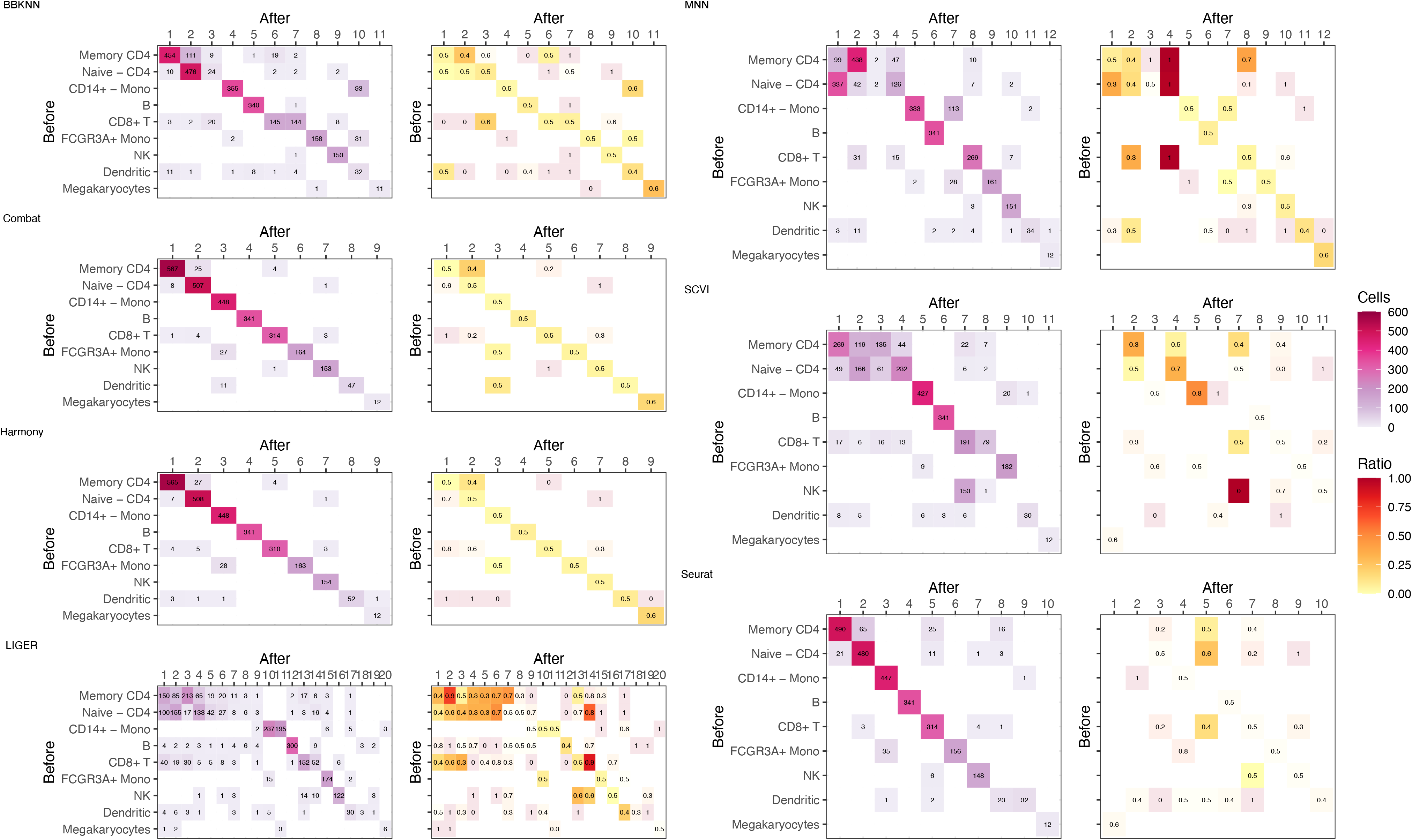
Confusion matrix of Leiden cell type clustering for the PBMC3K data. On the left the squares are colored by number of cells. On the right the coloring is by the ratio of cells in batch 1 with the color intensity altered by the number of cells in that square . On the y axis the cell types before batch correction and on the x are the cell types after. The best case result here is only values on the diagonal, where cells in a certain cell type cluster remain with the cells they shared a cluster with, after correction and for all ratios on the graph on the right to be 0.5.

We find that Harmony, Combat, and Seurat show the least amount of cluster confusion, and Harmony and Combat perform better on the imbalance between batches. BBKNN and MNN create more clusters, and some clusters show high imbalance, indicating an enrichment of batch effect creating a new cluster. On the other hand, SCVI and LIGER create the largest change in cluster number and composition with several of the clusters created being clear batch effects. Additional numerical evaluations of changes in cell type cluster can be found in the supplement.

### Differential expression

A common part of single cell RNA-seq workflow is finding those genes that are differentially expressed between some condition or state (Luecken and Theis 2019). One of the aims is to identify the marker genes that characterize clusters, that can then be used to identify one or more cell types that are represented in the cluster. We performed differential expression using MAST(Finak et al. 2015).

In order to estimate the effect of batch correction on gene expression we examined the change in the differential expression after applying each method, we estimated the degree of change using the number of differentially expressed genes between two cell type clusters, before and after batch correction. We performed this comparison using two models, one with only the cluster type as a covariate and then one model with cluster and batch as covariates in MAST.

We compare the methods using the PBMC3K data and the Mouse brain data. Additionally, to establish a positive control for this comparison, we simulate a true batch effect, by reducing the expression of the top 100 differentially expressed genes within a cluster for one of the batches.

For PBMC3K we found differentially expressed genes between B-cells and CD8+ T-cells, in the original dataset this resulted in 179 and 153 differentially expressed genes using cluster or cluster and batch as a covariated respectively (Figure 3A). MNN and Seurat report over 800 differentially expressed genes, indicating that the batch correction leaves a statistically significant trace on the gene expression that will result in too many false positives. COMBAT finds an unusal number of statistically significant genes when correcting for the batch covariate, this turns out to be driven by the fact that COMBAT performs batch correction independent of clusters or the KNN graph (Supplementary). Overall, SCVI finds a similar number of differentially expressed genes, but has a lower overlap with the results from original dataset. BBKNN, recovers 90% of the differentially expressed genes, but Harmony has the best performance overall.

**Figure 3:**
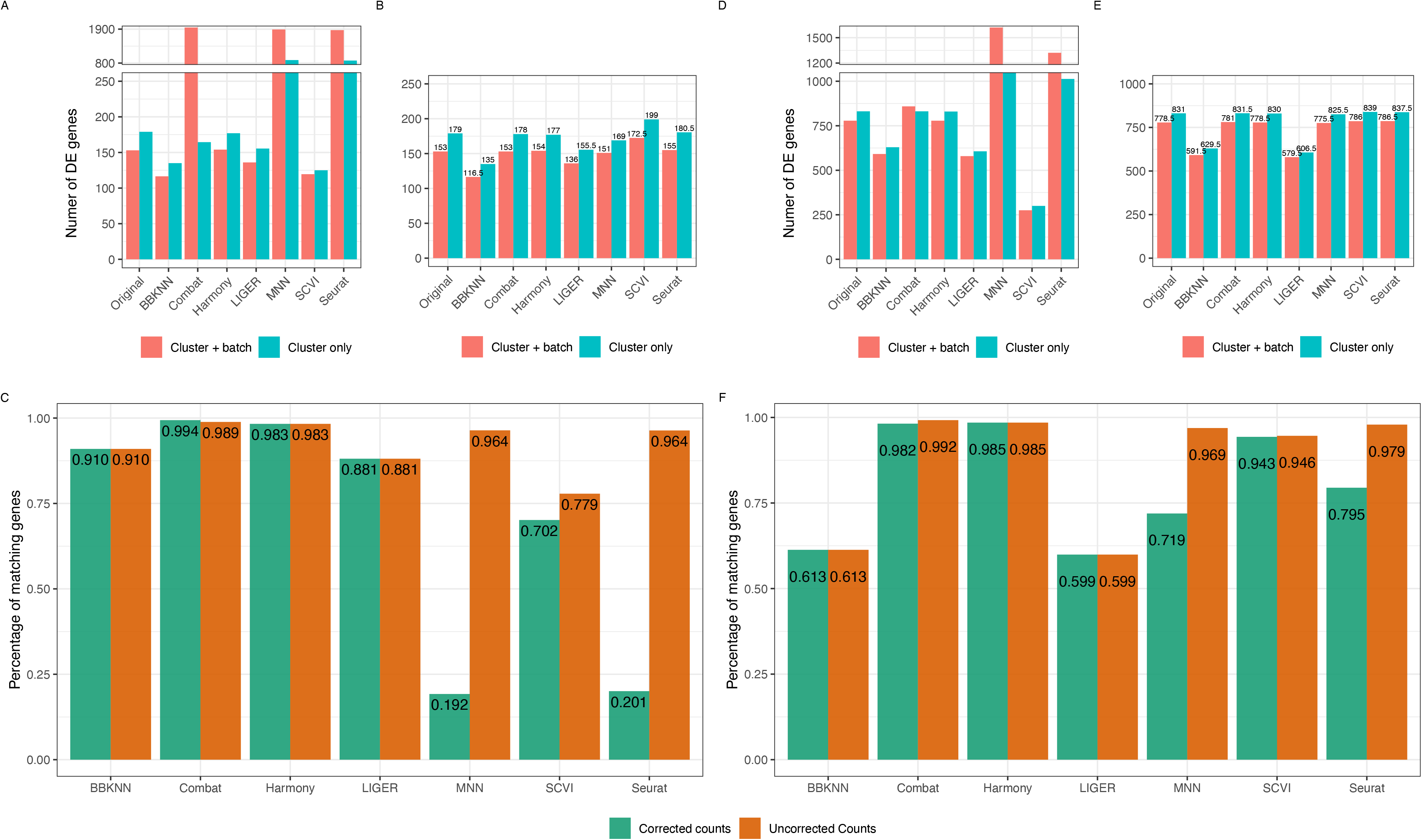
**A**: Number of statistically significant genes, out of the 2000 most variable, in the PBMC3K dataset. The best result here being those closest to the unaltered original data. **B**: Number of statistically significant genes, out of the 2000 most variable, in the PBMC3K dataset, when using the original uncorrected count data but with clusters calculated on the corrected count data. **C**: Percentage of the statistically different genes after correction in the PBMC3K dataset, that are also found in uncorrected data for the cluster only model. The color indicates which count matrix was used for modelling. For uncorrected counts the uncorrected count matrix was used for the model testing but with cell type clusters calculated on the corrected count matrix. A lower ratio indicates that the process of batch correction alters the genetic expression profiles of clusters. **D**,**E** and **F**, respectively, are the same but created using the Mouse brain dataset.

**Figure 4:**
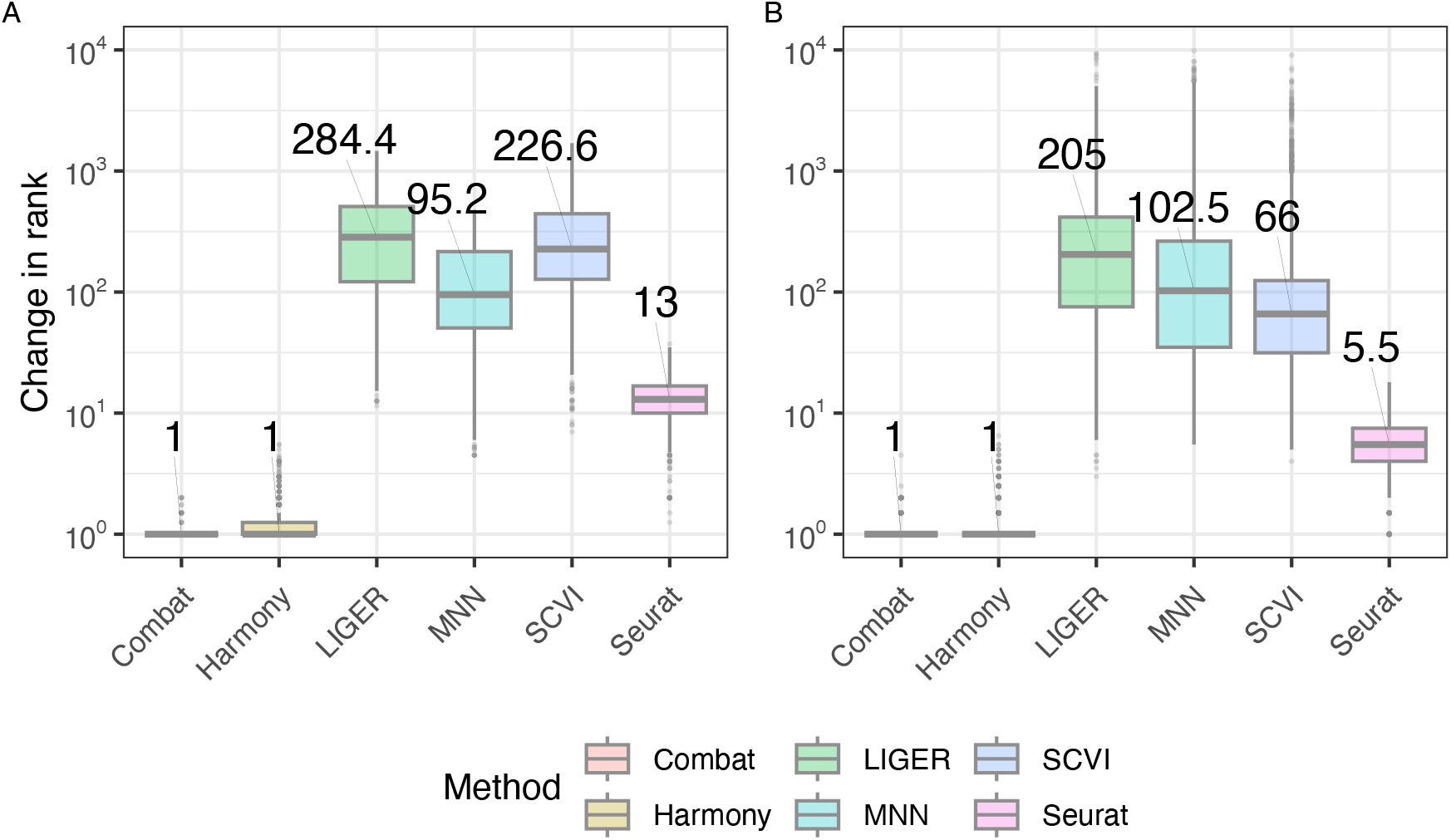
**A**:The PBMC3K data, **B**: Mouse brain data, Median change in absolute rank difference for the top 30 nearest neighbors of each cell for each method in the lower dimensional embedding, after batch correction.

**Figure 5:**
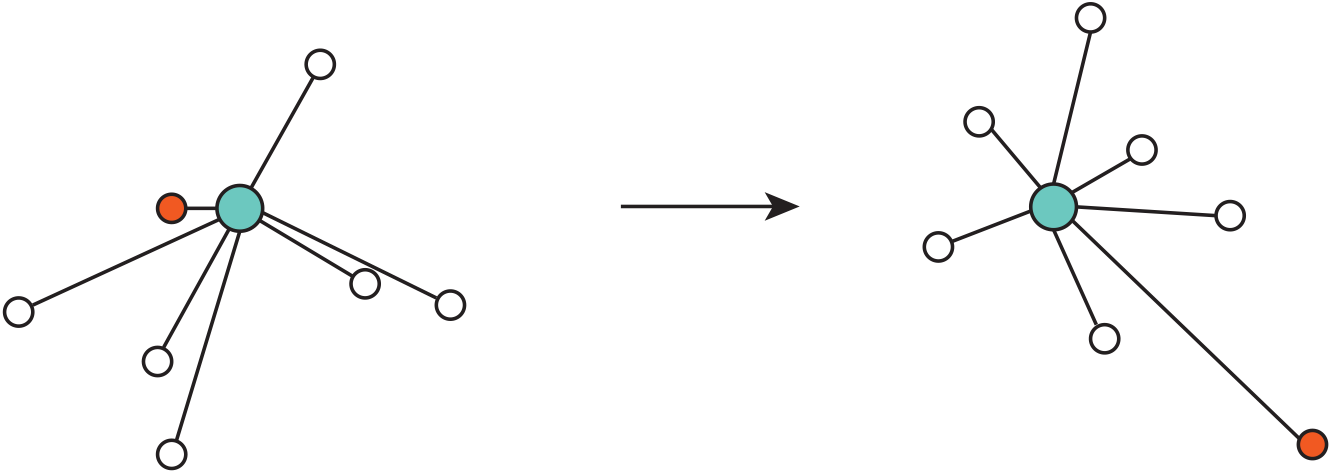
We see a NN graph of the center cell *c*, in blue, before and after batch correction. Before batch correction the orange cell has a rank of 1, i.e. it is the closest neighbor to cell *c*. After batch correction it has a rank of 7. We define rank displacement as this change in NN rank.

MNN, Seurat and SCVI perform better on the Mouse brain data compared to the PBMC3K data. BBKNN and LIGER perform worse. Similarly to the PBMC3K dataset, Seurat and MNN report a considerably higher number of differentially expressed genes compared to the other methods. For all the methods tested we see an improvement when using the clustering created on the uncorrected data.

To isolate whether the differences between the results on the original dataset and the batch corrected versions, we repeated the differential expression analyses, but used the uncorrected count matrices and the cluster assignments from the batch corrected datasets. Comparing panels A and B for the PBMC3K dataset and panels D and E for the Mouse brain dataset in Figure 3 we see that the poor performance of MNN, Seurat and COMBAT is driven by the modifications of the count matrix rather solely the cluster assignments, whereas Harmony, Liger and BBKNN do not adjust the count matrix.

We repeated the analysis on the PBMC3K and Mouse brain datasets with batch effects simulated an identified the same patterns as in the absence of batch effects (Supplement).

### Nearest neighbor rank – embeddings

The KNN graph is a fundamental object that is used as input in several widely used algorithms in single cell RNA-seq data analysis, e.g. graph based clustering (Leiden) and the creation of UMAP figures. Thus, we want to measure the extent of the change in the neighborhood composition of cells, created by the process of batch correction.

Comparing the rank displacement in the lower dimensional embedding, we see that, Combat and Harmony consistently have the smallest rank displacement, while SCVI, Liger and MNN have the highest displacement, representing on average no overlap between the top 30 neighbors.

These comparisons use dimensionality reduced data, top 50 PCA components for count-based methods (Combat, MNN, Seurat), corrected embedding (Harmony, Liger, SCVI), while BBKNN was left out since it alters the KNN graph, but not the distances required to order neighbors.

We observe a similar performance of the count-based methods, when the rank displacement is computed on the full count matrix, rather than a lower dimensional embedding (Supplement).

## Discussion

We tested seven methods for batch correction and by constructing randomized pseudo batches, we examined the degree of change, in various metrics, before and after batch correction.

Overall we see that the methods tend to fall in 3 categories of performance. Combat and Harmony perform the best or close to it in all the evaluations we perform. Those two methods introduce the fewest artifacts into the data during the process of batch correction. Seurat introduces artifacts that are more easily identifiable, but seems to retain the overall structure of data when comparing clusters. Finally, MNN, SCVI, LIGER and BBKNN, all introduce a significant change that alters local neighborhood structure, clusters and differential expression results.

While Combat performs well in the tests performed in this paper, it has been found to be outperformed by more modern methods when it comes to removing batch effects (Haghverdi et al. 2018) (Tran et al. 2020). Harmony is well suited for a variety of integration with a low amount of artifacts being introduced, in cases where there is little batch difference.

The role of batch correction in single cell RNA-seq analysis it twofold. First, we want account batch effect so that cells of the cell type are clustered together in the combined dataset. Second, after forming clusters, we need to account for the batch effect in other downstream methods, e.g. differential expression or finding marker genes.

The count-based methods, MNN, Seurat, and Combat, aim to solve both problems by altering the count matrix, whereas the other methods only tackle the first issue. Based on our results we can only recommend Harmony for general purposes since it performs the best all metrics. While Harmony does not correct the count matrix, the batch covariate can be used as input in statistical methods, e.g. MAST, to account for the batch effects.

In general, we would advise future method developers to focus solely on correcting local structure for batch effects and use classical methods to account for batch effects in statistical tests.

## Methods

### Simulated data

We simulated a difference in expression level for a particular cell type by performing ad hoc clustering and reducing the UMI counts by 50%, before any normalization or correction, for the top 100 most differentially expressed genes. The clusters modified were microglial cells in the mouse brain dataset and in B-cells in the PBMC3K dataset.

### Analysis workflow

For all the datasets we used Scanpy (Wolf et al. 2018) to perform basic preprocessing. In addition to filtering out cells with few detected genes, and genes detected in few cells, cells with high fraction of mitochondrial counts were also removed. Library size normalization was then performed before log-transformation. In the case of the mouse brain data, in addition to this the data was scaled and variation was regressed out.

To standardize the comparison the same 2000 highly variable genes, calculated on the uncorrected data, were used as input except in the case of the the alternative workflows of LIGER and Seurat, where we find the most variable genes using methods provided in the libraries themselves, as is recommended by the authors (Butler et al. 2018) (Welch et al. 2019). This alternative comparison is presented in the supplement.

A k-NN graph was created using the PCA embedding using Scanpy with default values. Following that cell clusters were calculated using the Leiden algorithm with default settings (Traag et al. 2019).

Each step in the workflow was run in Scanpy or a python library if possible. For those batch correction methods that only provide libraries in R, the preprocessing was performed in python then the batch correction was run in R using Seurat where possible. All the datasets were kept in the Anndata format in python (Virshup et al. 2023). The data was exported to files and imported in R when required.

### Downsampling

To estimate a baseline of technical variation to expect when integrating datasets with different modalities we compare our results to simulated batch effects with downsampling. Our aim is to provide a lower bound of change in our evaluation criteria. We want to evaluate if the batch correction methods alter the data more than simply downsampling one batch and using the combined data without additional correction. For the Heart and PBMC4K Fastq files, we downsampled the data using BUStools (Melsted et al. 2019).For both datasets we downsampled the reads of all cells within one batch to 50%. To standardize these results we used only cells found in one of the pseudo batches in the original uncorrected dataset. There is therefore no cell filtering performed on the Anndata file created using the downsampled BUS files.

For each method there exists a plethora of different configurations and parameters that can be configured to adapt the method to different scenarios. In this case the default parameters were used whenever possible. In some cases we selected parameters to enable easier comparison between the different methods, e.g. selecting the top 2000 most variable genes instead of using the default method in Scanpy. This ensured that for each method the number of highly variable genes was the same. Additionally, whenever possible we provided a seed to make a randomized method return deterministic results.

## Evaluation criteria

### Change in nearest neighbor rank

Neighborhood structure is an important part of downstream analysis in scRNA-seq. The neighborhood of different cells allows us to get an understanding of the local structure. Commonly a k-NN graph is created to define the k nearest neighbors of each cell for all cells. An important downstream result that depends on the k-nn graph is the cluster analysis. The commonly used leiden (Traag et al., 2019) method uses the NN graph to perform clustering. In each case the k-NN graph was calculated using the default parameters in Scanpy.

In order to measure the effect that the batch correction methods have on the nearest neighbor structure we examine the location or rank, in terms of order of closest to furthest, of nearest neighbors of each cell before and after batch-correction. For those methods that change the count matrix we use the log normalized count matrix to calculate the change in NN rank. The neighborhood structure is commonly determined in some lower dimensional space. In order to get a concise measure of the extent of the nearest neighbor displacement we additionally, examine the change in nearest neighbor structure in the lower dimensional embedding. We compare the NN rank on the PCA embedding on the uncorrected data to a PCA embedding created on the corrected count matrices, except in the case of Harmony, LIGER and SCVI, which return a corrected embedding that is used for comparison.

We calculate the following for each cell *i* and nearest neighbor(NN) *j*.

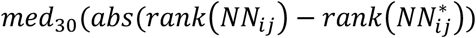

Where *med*_30_ is the median over the top 30 nearest neighbors and 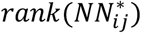 is the rank of the j-th NN of cell i after batch correction. Differential expression

In order to assess if batch correction methods have an effect on differential expression results we compare the number of statistically different genes between 2 distinct cell clusters in the data, before and after correction. As we only tested the top 2000 most variable genes, the maximum number will be 2000. For the PBMC3K dataset we compared clusters that represented B-cells and CD8+ T-cells. For the mouse brain data we considered smooth muscle cells and microglial cells. These cell types were chosen because they have highly differentially expressed, unique markers (Tasic et al. 2016) (Li et al. 2022).

We used 2 different MAST models and each model is then run for each gene. We use as input into the model, only on the cells in the clusters being compared. The MAST model is a two-part hurdle model that models the rate and level of expression for cells for each gene. Which allows for the controlling of covariates. We included batch and cluster as covariates to assess the statistical significance of each on gene expression. The models we implemented are as follows:

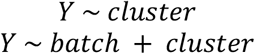

A likelihood-ratio test(LRT) is then performed to assess the significance of the covariate in question. The cluster covariate in the cluster only model, and then both covariates in the batch and cluster model. An LRT is performed for each gene.

We used as a ground truth, the genes determined to be differentially expressed in the original dataset without any batch correction. For each batch correction method we determined the differentially expressed genes between the two clusters. The clusters used the analysis were chosen as the two clusters that have the same particular marker genes as the clusters in the original dataset. We also performed the differential expression analysis using the exact set of cells as in the analysis without batch correction. Finally, we repeated the differential expression analysis using the cluster assignment from batch corrected data, but the original count matrix as input to MAST.

As an additional comparison of the effect of correcting the original count matrix, we run the same models using MAST, but instead of using the corrected count matrix we use the corrected cluster assignments provided by the methods or calculated after using the methods, but the count matrix contains the uncorrected original counts.

## Data access

Publicly available datasets from 10X Genomics were used for analysis and simulations.The PBMC3K data is derived from Peripheral blood mononuclear cells from a healthy donor, generated by 10x Genomics, with a sequencing depth of 69000 reads per cell. The Mouse brain dataset is derived from brain tissue of two E18 mice, with around 18500 reads per cell. We use a 20k cell sample provided by 10x Genomics. The PBMC4K dataset is similar to the PBMC3K dataset except the sequencing depth is higher at around 87000 reads per cell. The Mouse heart data was derived from the whole heart of a mouse, with a sequencing depth of around 83000 reads per cell. All code used to perform the analysis is available at https://github.com/pmelsted/AM_2024

## Acknowledgements

S.E.A. was supported by the University of Iceland Doctoral Grants and the Icelandic Research Fund Project (218111-051). The authors would like to thank Ali Sina Booeshaghi for feedback on the manuscript.

