## Supplementary Material for "Batch correction methods used in single cell RNA-sequencing analyses are often poorly calibrated"

#### Additional NN displacement comparisons

We evaluated the methods on four datasets, by comparing the nearest neighborhood (NN) structure before and after batch correction. Briefly, the rank displacement measures how far back in line a neighbor of a cell moved after correcting the data (Methods), a high value for this rank displacement metric would indicate that a method alters the NN structure for a large number of cells.

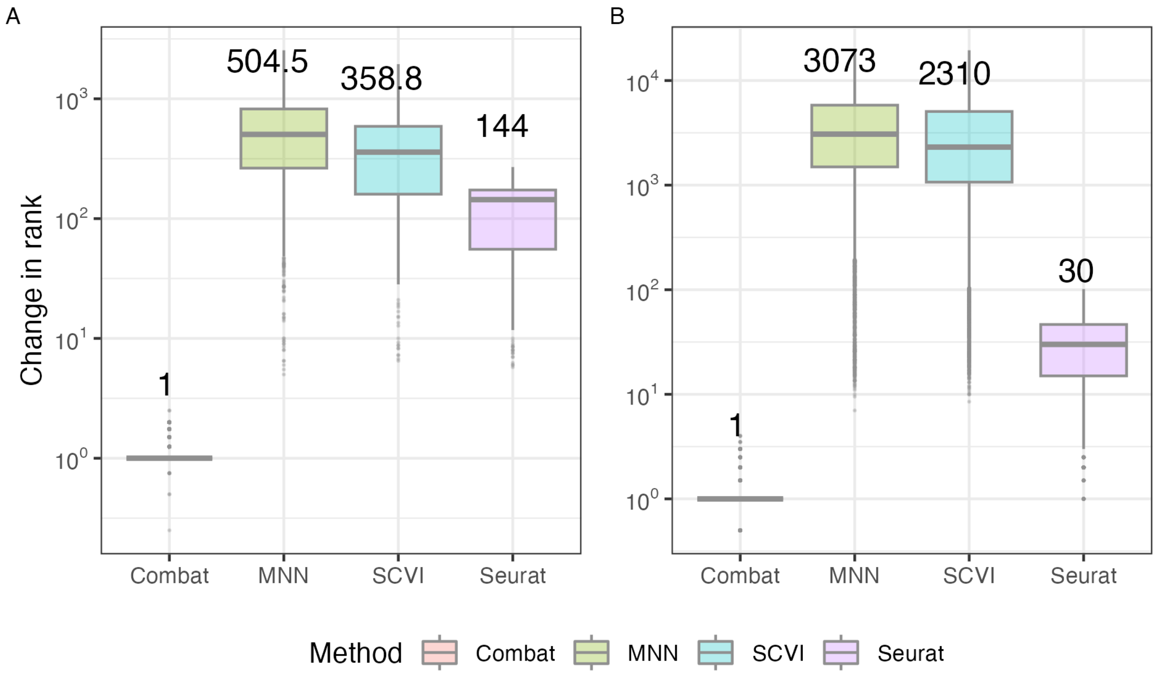


**Supplementary figure 1**: A:The PBMC3K data, B: Mouse brain data, Showing median absolute NN rank displacement for the top 30 NN of each cell, after applying batch correction.

Combat performs best when using this metric, the internal order of neighbors is very well preserved. When taking into consideration the total number of cells in each dataset, around 2700 and 20000 cells for the PBMC3K and mouse brain data respectively, MNN and SCVI cause a high degree of change in the neighborhood structure of cells.

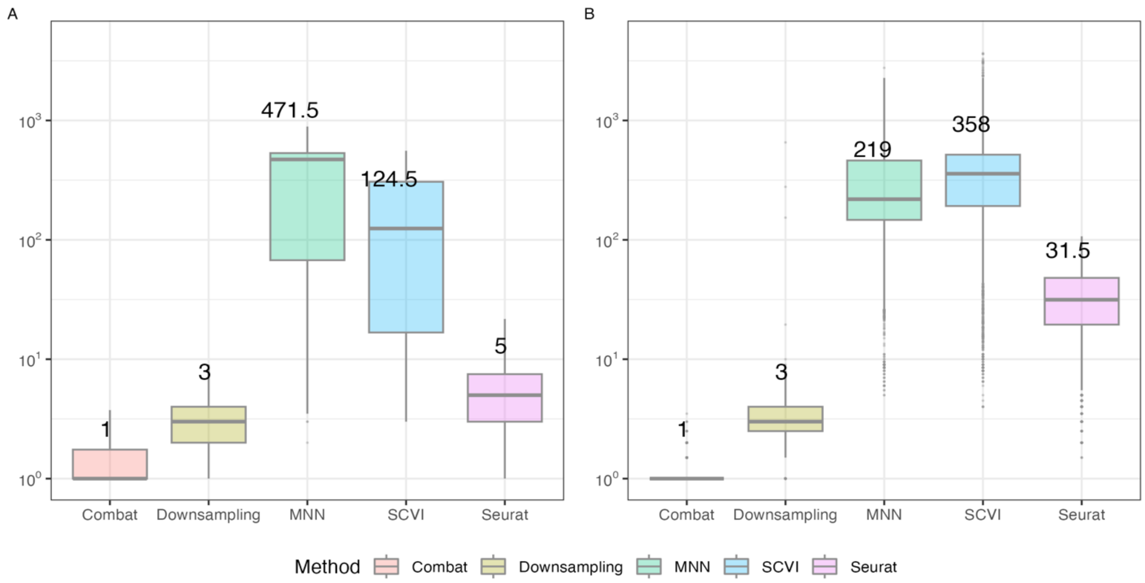


**Supplementary Figure 2**:The Mouse heart data, B: PBMC4K data, Showing median absolute NN rank displacement for the top 30 NN of each cell, after applying batch correction. With the added downsampling.

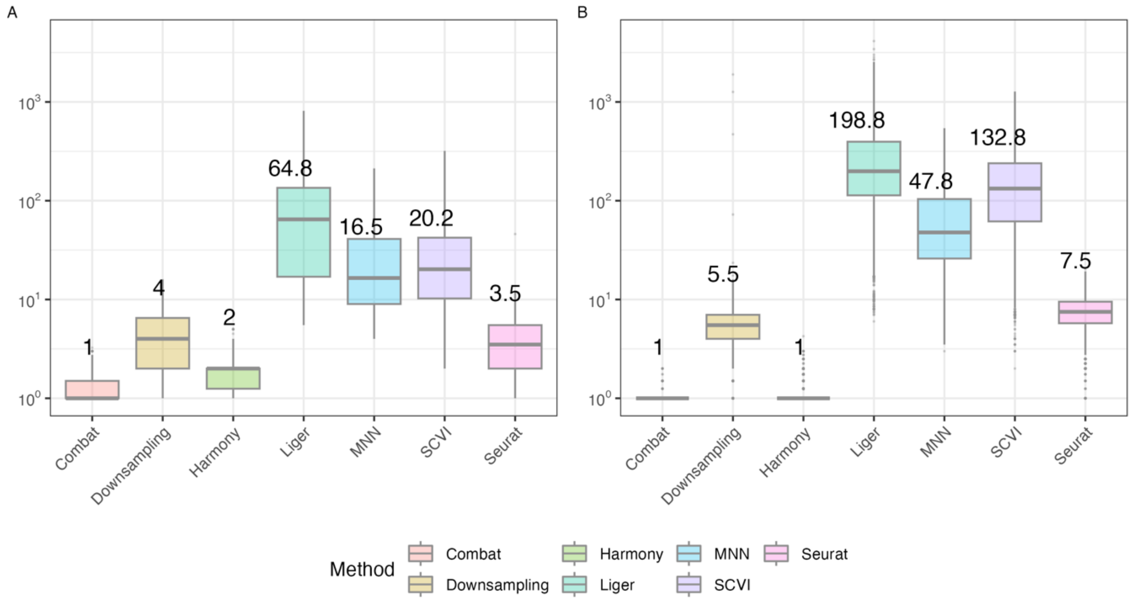


**Supplementary Figure 3**: A: Mouse heart B: PBMC4K Median change in absolute rank difference for the top 30 nearest neighbors of each cell for each method in the lower dimensional embedding.

### Changes in cell type clusters - Mouse brain

We repeat the cell confusion matrix tables with the mouse brain data, with the y axis being cluster assignment before batch correction and x axis before. The columns have been ordered in such a way that we get similar cell numbers on the diagonal. The number of cells and cell type clusters increase but see similar performance as we did with the PBMC data. With Combat and Harmony altering the cluster structure the least.

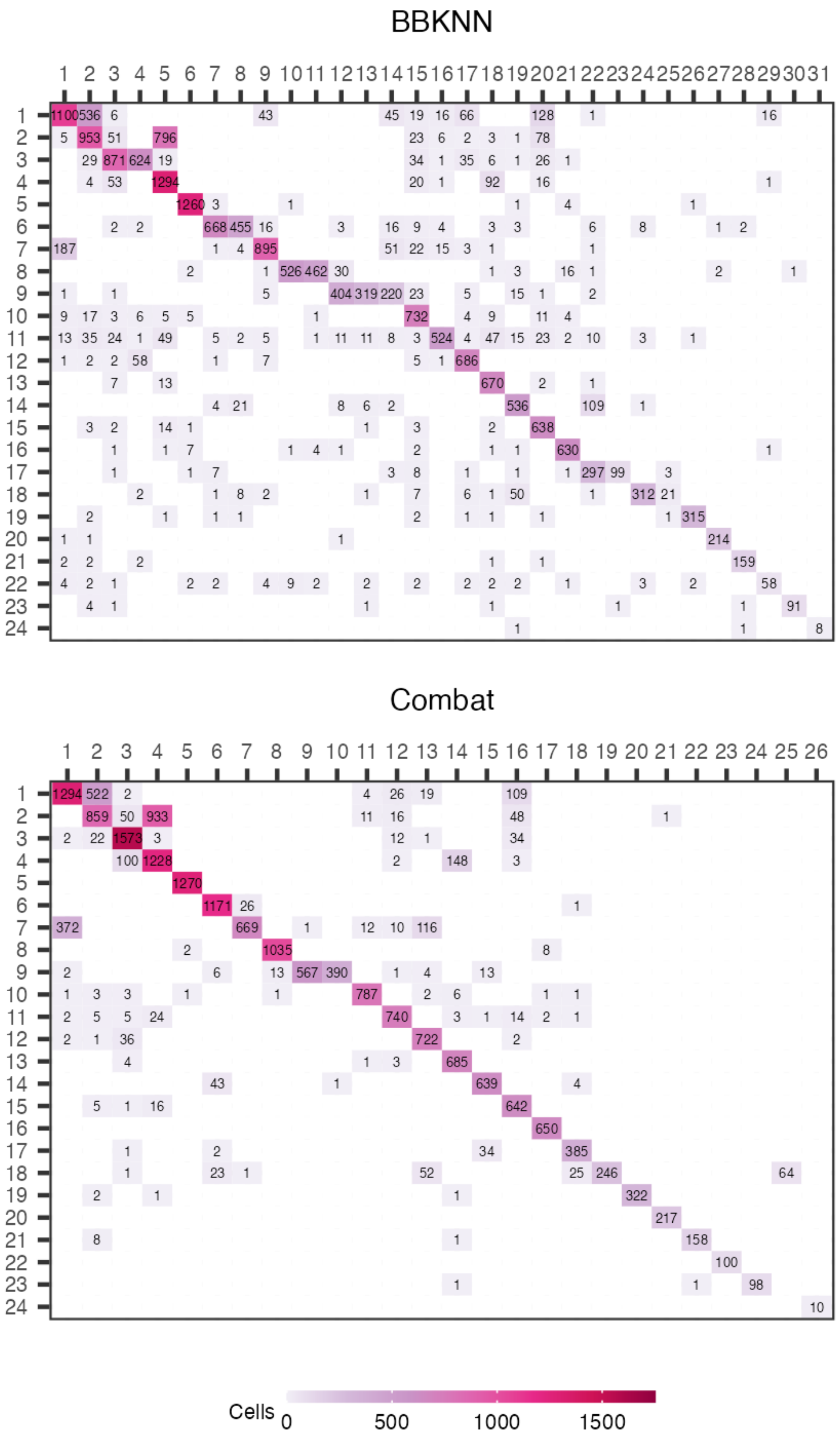


**Supplementary figure 4**: Confusion matrix for cluster assignment, before and after batch correction for the Mouse brain data using BBKNN and Combat.

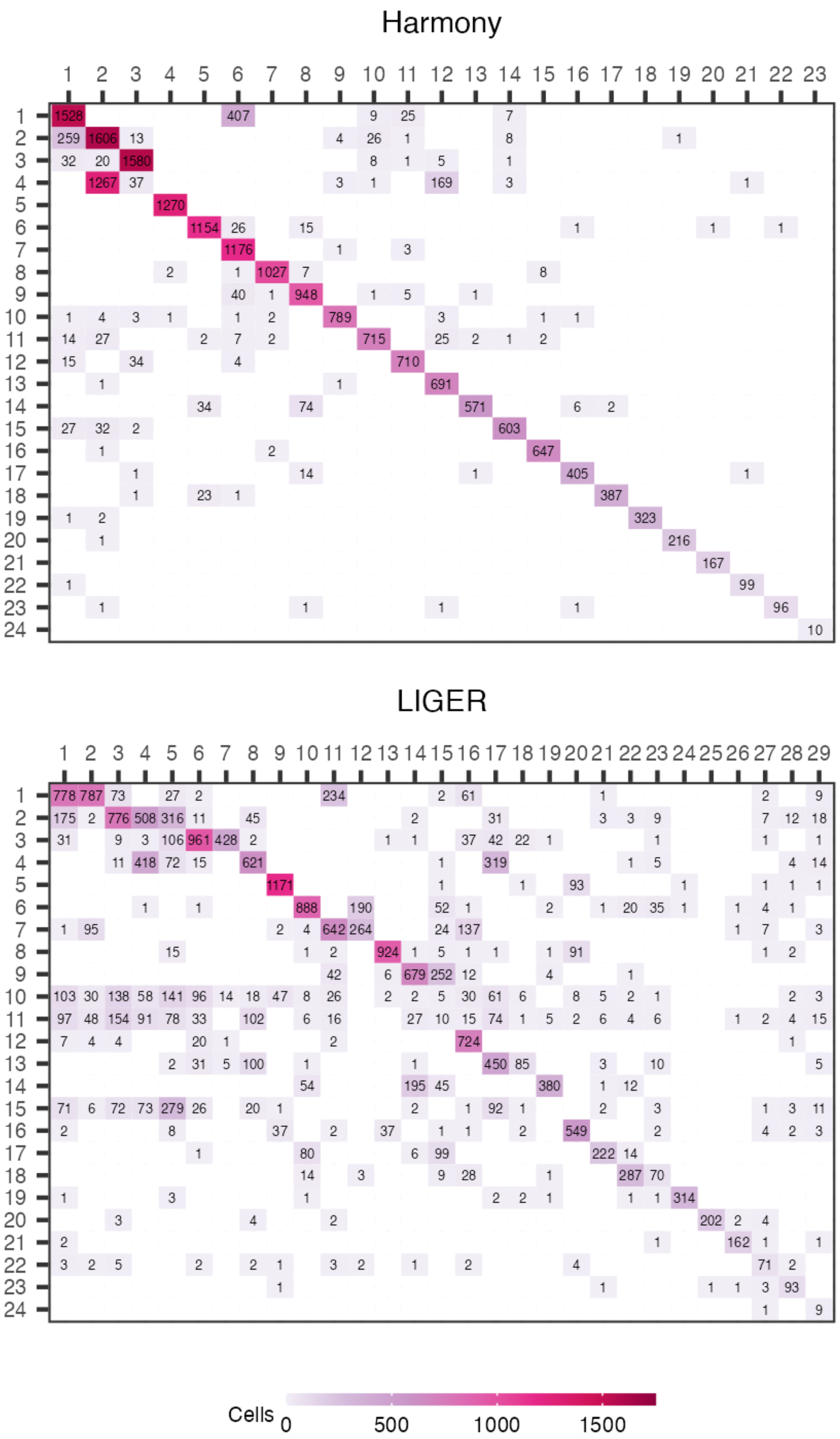


**Supplementary figure 5**: Confusion matrix for cluster assignment, before and after batch correction for the Mouse brain data using Harmony and LIGER.

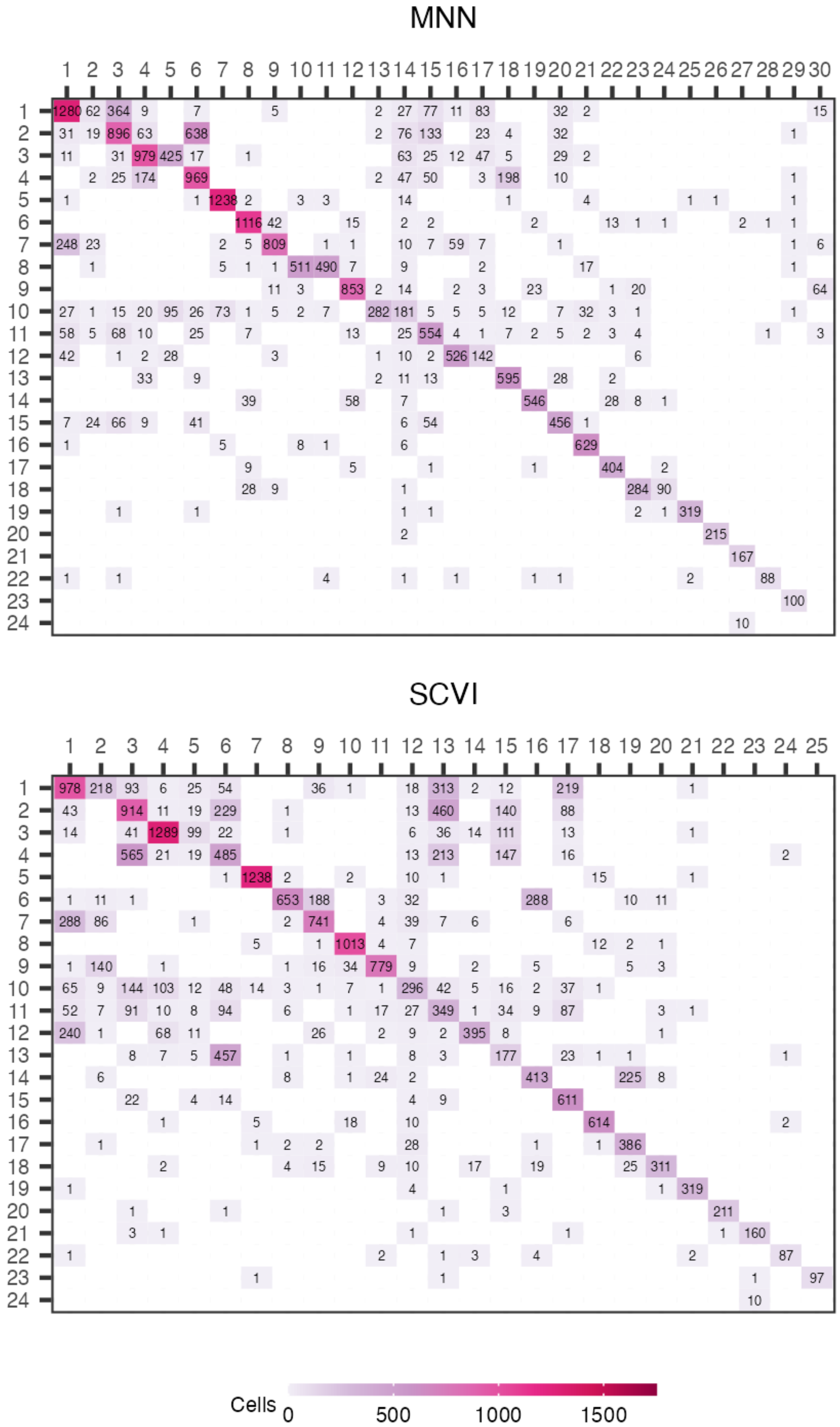


**Supplementary figure 6**: Confusion matrix for cluster assignment, before and after batch correction for the Mouse brain data using MNN and SCVI.

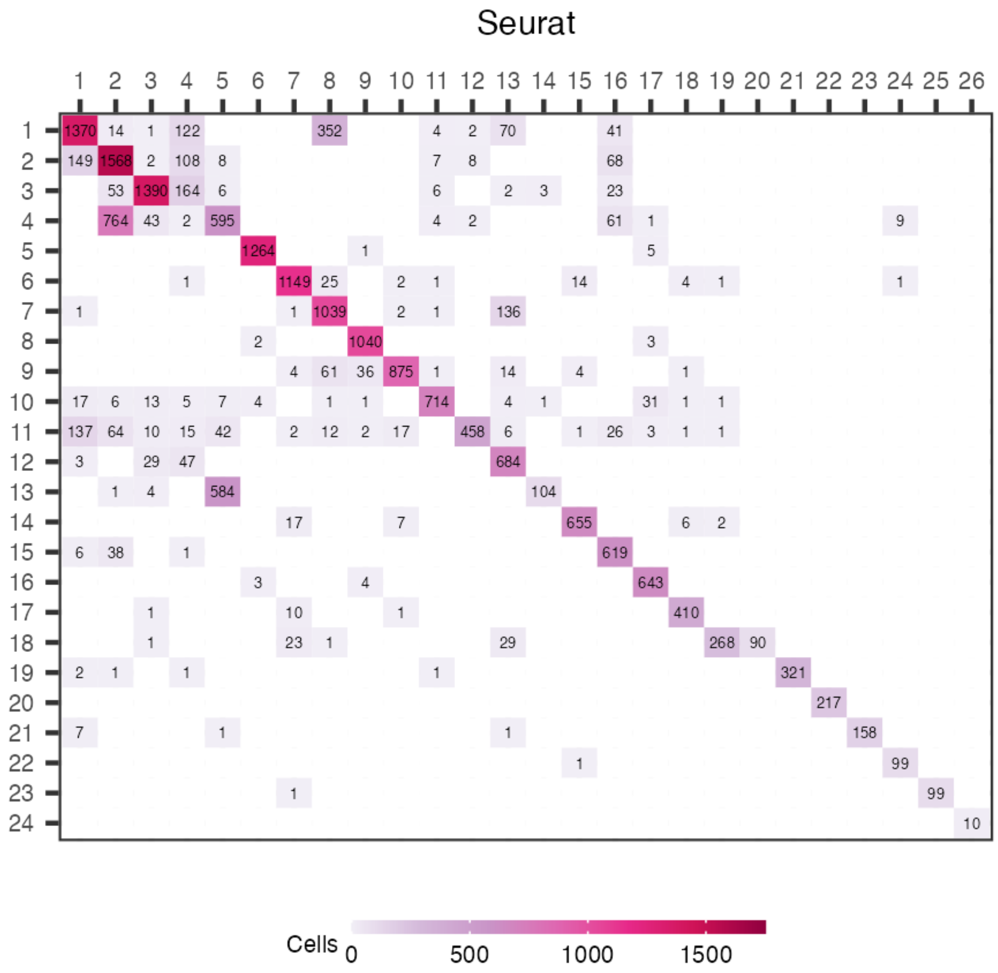


**Supplementary figure 7**: Confusion matrix for cluster assignment, before and after batch correction for the Mouse brain data using Seurat.

### Consensus cluster metric

In order to assess the effects of batch correction on the clustering, we compared the original clusters to the clusters created on the batch corrected data. While we provide the confusion matrices in the results, a concrete numerical value is provided here that should provide comprable results. Because the number of clusters can differ before and after batch correction and because this is a multiclass problem, more traditional classification scores cannot be used in order to evaluate the change. Almost all the cells of one cluster going into another and vice versa is not a negative thing in this context. A single cluster going into multiple or multiple clusters merging into a single one is a negative outcome. To assess this change we devised a metric that ranges from 0 to 1 that aims to evaluate the degree of change the clustering undergoes after applying batch correction methods. In this case 0 is no change to the clusters and a value of 1 indicates that the clustering has been altered considerably.

If we define $c_{i,j}$ to be the number of cells in cluster i before batch correction and cluster j after. $C_i = \sum_j c_{i,j}$ is the number of cells in the i-th cluster before batch correction and $C_j^* = \sum_i c_{i,j}$ is the number of cells in the j-th cluster after batch correction. We can then define the following.

| Confusion matrix | | | |
|---|---|---|---|
| Cluster | 1 | ... | m |
| 1 | $c_{1,1}$ | ... | $c_{1,m}$ |
| ⋮ | ⋮ | | ⋮ |
| n | $c_{n,1}$ | ... | $c_{n,m}$ |

$$T_i = \frac{C_i - \max(c_{i,1}, \dots, c_{i,m})}{C_i}$$

$$T_j^* = \frac{C_j^* - \max(c_{1,j}, \dots, c_{n,j})}{C_j^*}$$

Then we define $\mu_1$ and $\mu_2$ as the weighted mean of $C_i$ and $C_j$ respectively:

$$\mu_1 = \frac{\sum_i T_i w_{C_i}}{\sum_i w_{C_i}}$$

Where $w_{C_i} = \frac{\sum_j c_{i,j}}{\sum_{i,j} c_{i,j}}$ is the number of cells in cluster i, divided by the the total number of cells. Similarly we define

$$\mu_2 = \frac{\sum_j T_j^* w_{C_j}}{\sum_j w_{C_j}}$$

Where $w_{C_j} = \frac{\sum_i c_{i,j}}{\sum_{i,j} c_{i,j}}$

Then we get the consensus cluster value as the mean between $\mu_1$ and $\mu_2$

$$CC = \frac{\mu_1 + \mu_2}{2}$$

| Data | BBKNN | Combat | Harmony | LIGER | MNN | SCVI | Seurat |
|---|---|---|---|---|---|---|---|
| **PBMC3K** | 0.13 | 0.04 | 0.04 | 0.35 | 0.17 | 0.24 | 0.09 |
| **Mouse brain** | 0.24 | 0.11 | 0.12 | 0.39 | 0.26 | 0.32 | 0.15 |
| **PBMC3K Simul** | 0.16 | 0.06 | 0.05 | 0.36 | 0.22 | 0.27 | 0.08 |
| **Mouse brain Simul** | 0.28 | 0.13 | 0.13 | 0.38 | 0.28 | 0.33 | 0.23 |

**Supplementary table 1:** Consensus cluster metric score after batch correction for 4 different datasets.

Methods with a lower score alter the clustering less. We see that Combat and Harmony achieve the lowest score over all the datasets. Seurat has a slightly higher ratio, with them performing quite differently between datasets. The three methods that have the highest degree of change in cell clusters after batch-correction are BBKNN, LIGER and SCVI.

| Data | BBKNN | Combat | Harmony | LIGER | MNN | SCVI | Seurat | Downsample |
|---|---|---|---|---|---|---|---|---|
| **PBMC4K** | 0.17 | 0.02 | 0.03 | 0.31 | 0.19 | 0.18 | 0.08 | 0.05 |
| **Heart** | 0.15 | 0.07 | 0.09 | 0.32 | 0.18 | 0.19 | 0.11 | 0.11 |

**Supplementary table 2**: Consensus cluster metric score after batch correction, including downsampling for the PBMC4K and Heart data.

We see  similar results here as in Supplementary table 1, with Harmony and Combat altering the clustering the least. By establishing a sort of baseline of results with downsampling we see that the Seurat methods alter the clustering similarly to downsampling one batch to 50% of reads. With MNN and SCVI causing the most change in the clustering, followed by BBKNN and LIGER.

### Seurat and Liger with original preprocessing

We attempted, within reason, to follow the recommended workflow for each method. To be able to compare the results of the differential expression we calculated the top 2000 most variable genes with Scanpy and used those genes as input into the data whenever possible. We include here the recommended workflow for Seurat and LIGER using only the genes selected using the provided methods in the respective libraries. We present a comparison of the results created using the genes we select and the native(alternative) workflow. While using different genes causes a slight difference in the metric, the order in which the methods perform, in the metrics we examine, does not change.

Nearest neighbour embedding

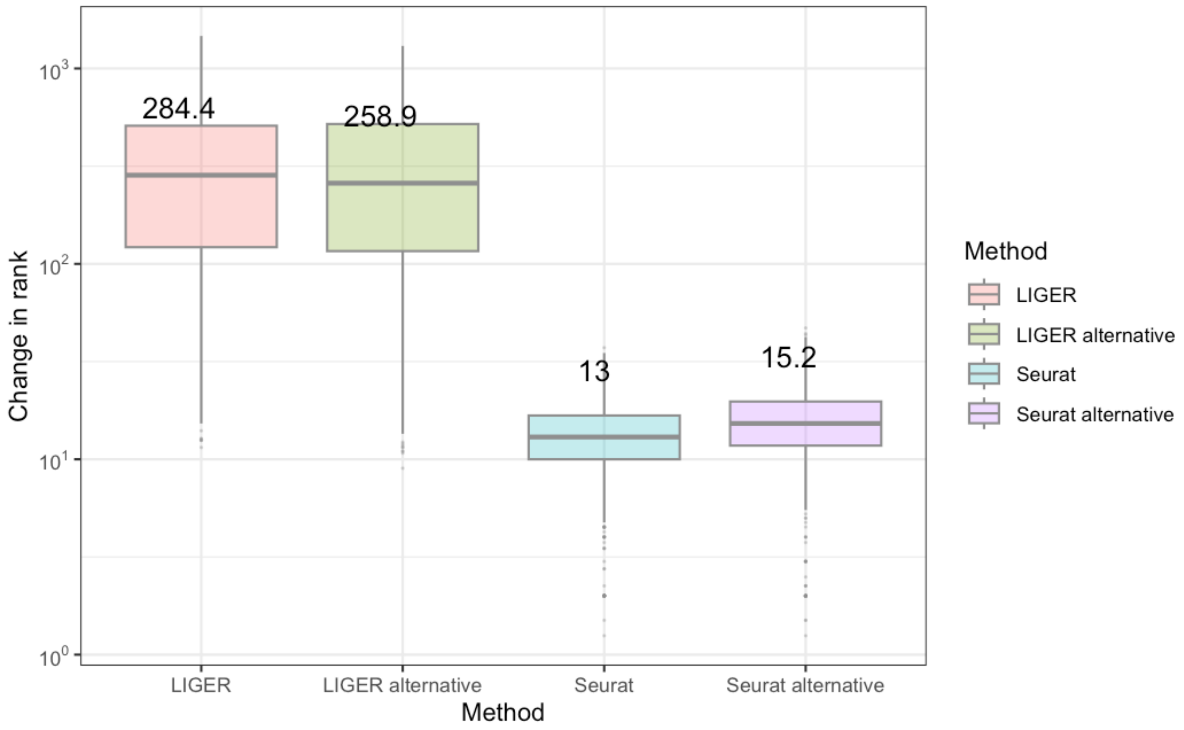


**Supplementary figure 8**: Change in NN rank before and after batch correction. Comparing the alternative default workflow for LIGER and Seurat with the one performed in this paper.

Cell confusion matrix – PBCM

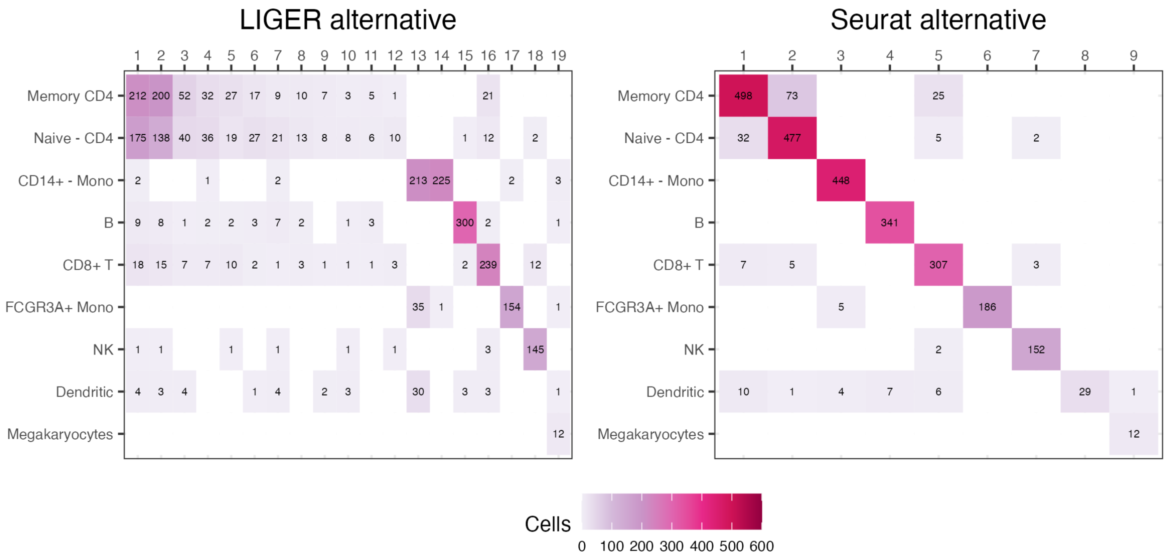


**Supplementary figure 9**: Change cluster confusion matrix before and after batch correction. Comparing the alternative default workflow for LIGER and Seurat with the one performed in this paper.

### Simulated differential expression results

#### PBMC3K simulated

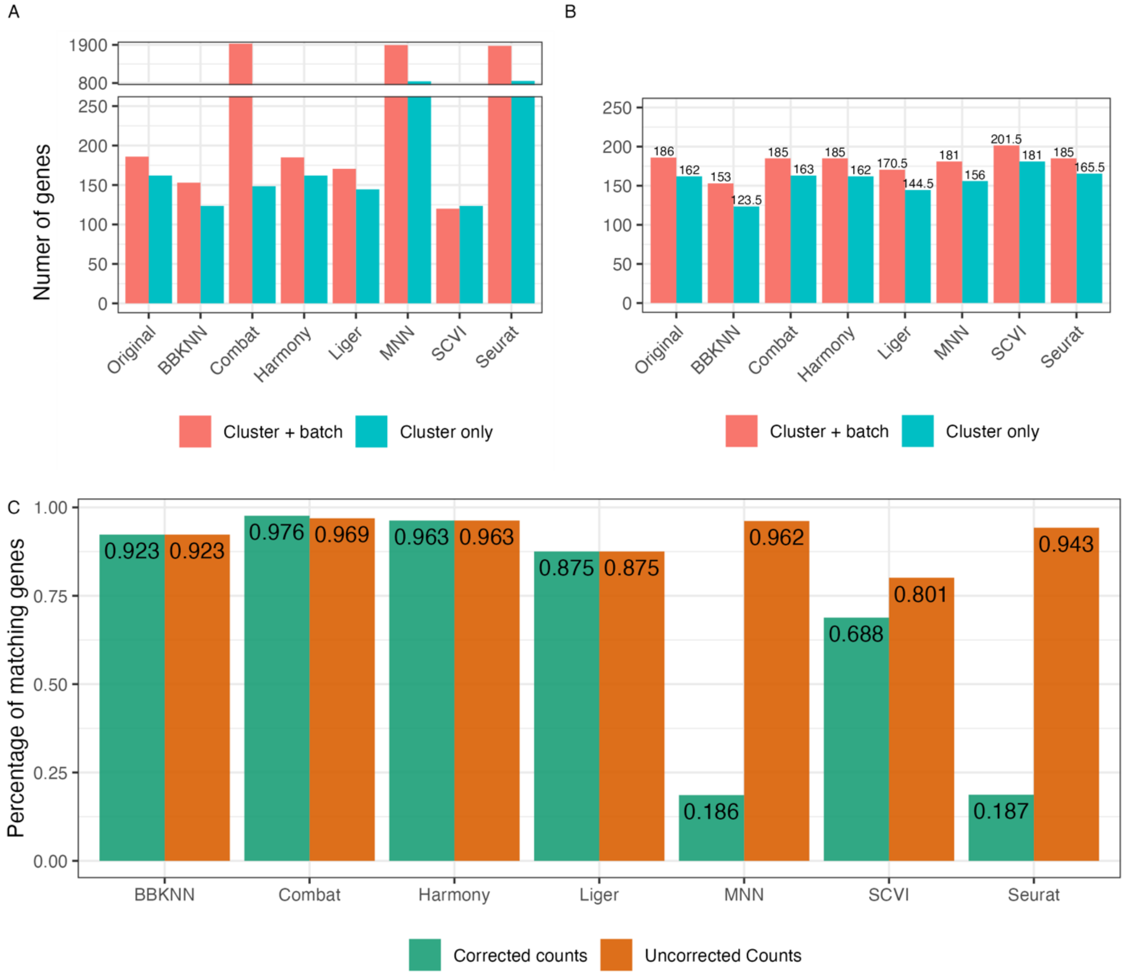


**Supplementary figure 10**: A: Number of statistically significant genes, out of the 2000 most variable. The best result here being those closest to the unaltered original data. B: Number of statistically significant genes, out of the 2000 most variable, when using the cell type clusters calculated on the original data. C: Ratio of the statistically different genes after correction,that are also found in uncorrected data for the cluster only model. The color indicates whether which count matrix was used for modelling. For uncorrected counts the uncorrected count matrix was used for the model testing but with cell type clusters calculated on the corrected count matrix. A higher ratio indicates that the process of batch correction alters the genetic expression profiles of clusters less.

#### Mouse brain - Simulated

As before we calculate the same metrics but on data where we have reduced the values for one batch, for the top genes in cells identified as Microglial cells.

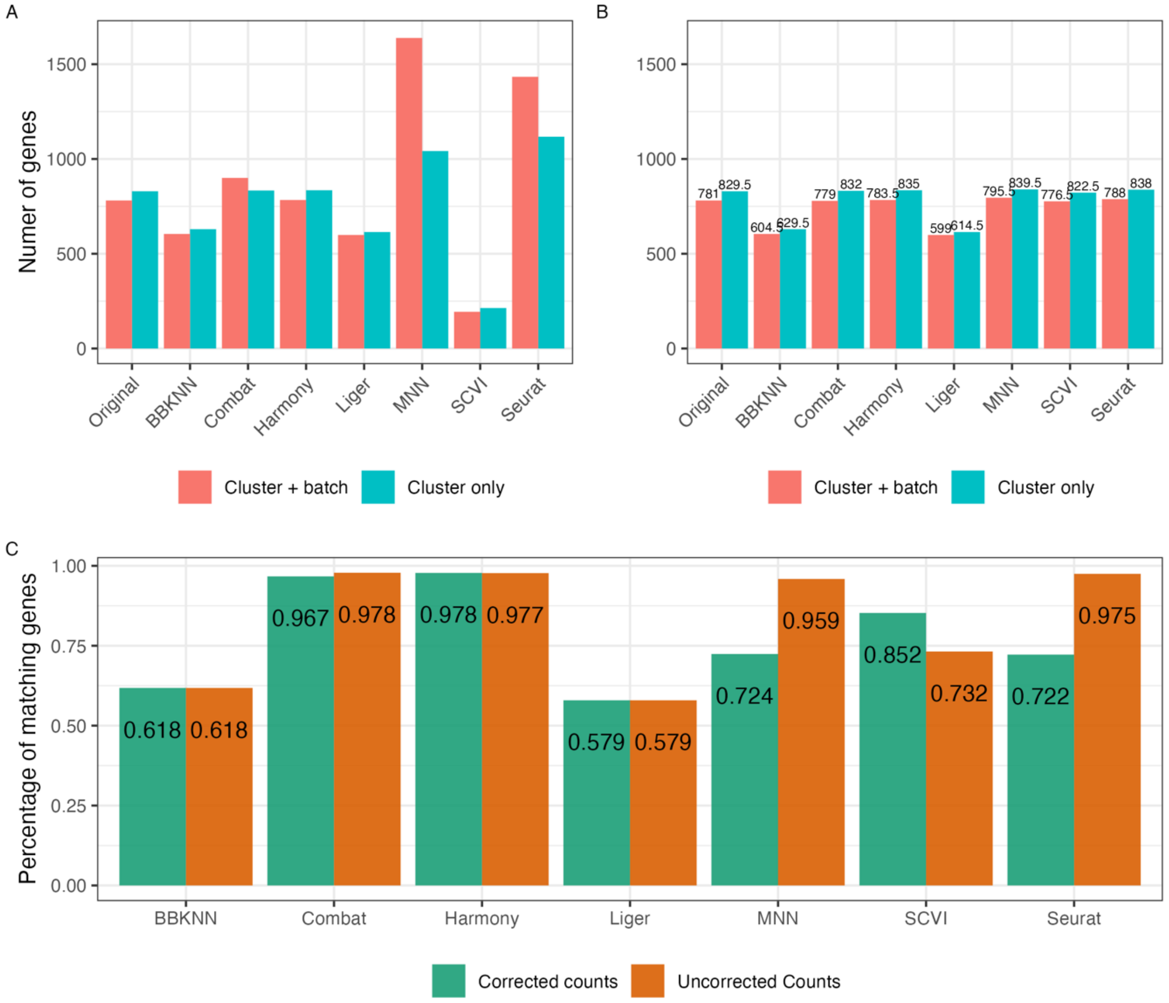


**Supplementary figure 11**: A: Number of statistically significant genes, out of the 2000 most variable. The best result here being those closest to the unaltered original data. B: Number of statistically significant genes, out of the 2000 most variable, when using the cell type clusters calculated on the original data. C: Ratio of the statistically different genes after correction,that are also found in uncorrected data for the cluster only model. The color indicates whether which count matrix was used for modelling. For uncorrected counts the uncorrected count matrix was used for the model testing but with cell type clusters calculated on the corrected count matrix. A higher ratio indicates that the process of batch correction alters the genetic expression profiles of clusters less.
